# Quantitative environmental DNA metabarcoding shows high potential as a novel approach to quantitatively assess fish community

**DOI:** 10.1101/2022.04.27.489619

**Authors:** Satsuki Tsuji, Ryutei Inui, Ryohei Nakao, Seiji Miyazono, Minoru Saito, Takanori Kono, Yoshihisa Akamatsu

**Affiliations:** Graduate School of Science, Kyoto University, Kitashirakawa-Oiwakecho, Sakyo-ku, Kyoto 606–8502, Japan; Graduate School of Science and Technology for Innovation, Yamaguchi University, 2-16-1 Tokiwadai, Ube, Yamaguchi, 755–8611, Japan; Faculty of Socio-Environmental Studies, Fukuoka Institute of Technology, Wajiro-higashi, Higashi-ku, Fukuoka 811–0295, Japan; Fisheries Division, Japan International Research Center for Agricultural Sciences, 1-1, Ohwashi, Tsukuba, Ibaraki 305–8686, Japan; Aqua Restoration Research Center, Public Works Research Institute, National Research and Development Agency, Kawashima, Kasada-machi, Kakamigahara, Gifu 501–6021, Japan

**Keywords:** Environmental DNA, Quantitative eDNA metabarcoding, qMiSeq, MiFish, Electrical shocker, Fish community

## Abstract

The simultaneous conservation of species richness and evenness is important to effectively reduce biodiversity loss and keep ecosystem health. Environmental DNA (eDNA) metabarcoding has been used as a powerful tool for identifying community composition, but it does not necessarily provide quantitative information due to several methodological limitations. Thus, the quantification of eDNA through metabarcoding is an important frontier of eDNA-based biomonitoring. Particularly, the qMiSeq approach has recently been developed as a quantitative metabarcoding method and has attracted much attention due to its usefulness. The aim here was to evaluate the performance of the qMiSeq approach as a quantitative monitoring tool for fish communities by comparing the quantified eDNA concentrations with the results of fish capture surveys. The eDNA water sampling and the capture surveys using the electrical shocker were conducted at a total of 21 sites in four rivers in Japan. As a result, we found significant positive relationships between eDNA concentrations of each species quantified by qMiSeq and both the abundance and biomass of each captured taxon at each site. Furthermore, for seven out of eleven taxa, a significant positive relationship was observed between quantified DNA concentrations by sample and the abundance and/or biomass. In total, our results demonstrated that eDNA metabarcoding with the qMiSeq approach is a suitable and useful tool for quantitative monitoring of fish communities. Due to the simplicity of the eDNA analysis, the eDNA metabarcoding with qMiSeq approach would promote further growth of quantitative monitoring of biodiversity.

**Highlights:**

- eDNA analysis and capture surveys of fishes were conducted at a total of 21 sites.
- Quantitative eDNA metabarcoding (qMiSeq) successfully quantified the eDNA of fishes.
- For each site, the estimated eDNA conc. reflected the abundance and biomass of fishes.
- For each taxon, the estimated eDNA conc. were comparable among sites.
- qMiSeq is a suitable and useful tool for quantitative monitoring of fish community.

## Introduction

Conspicuous loss in biological diversity is a current and fast-emerging global problem resulting in changes of ecosystem functions (Cardinale et al., 2012; Dornelas et al., 2014; Magurran et al., 2018). To effectively reduce biodiversity loss, a number of previous studies have argued for the importance of assessing and conserving ecosystem health using species abundance, evenness and richness as indicators (Blowes et al., 2022; Crowder et al., 2012; Hillebrand et al., 2008). Accordingly, researchers and resource managers have made efforts to quantitatively estimate biodiversity based on conventional survey methods, such as direct capture and visual census (Masuda et al., 2010; Miyazono et al., 2015; Yonekura et al., 2004). While these survey approaches provide us with valuable data, they also require the large amount of effort, time and expertise, which limits the feasibility and continuity of the research itself and the reliability of the data (Evans et al., 2017; Miya et al., 2020; Oka et al., 2021; Thomsen et al., 2012). Additionally, especially for endangered species, sampling activities in direct capture surveys may damage target-species populations and/or their habitats (Pimm et al., 2015; Rourke et al., 2022; Tsuji et al., 2018). To overcome these difficulties in the traditional survey methods, new approaches for accurate and effective biodiversity monitoring are being explored (Kissling et al., 2018; Rodríguez-Ezpeleta et al., 2021).

Environmental DNA (eDNA) analysis has been rapidly developed over the past decade as an alternative and/or complementary biomonitoring tool and has become widely used for organisms from various taxonomic groups (Boivin-Delisle et al., 2021; Deiner et al., 2017; Doi et al., 2021; Kelly, 2016; Kumar et al., 2020). Environmental DNA analysis enables us to estimate the presence of organisms via only the collection and detection of cellular materials shed from them in environments, including soil, water, and air (Ficetola et al., 2008; Kuwae et al., 2020; Lynggaard et al., 2022). The success and high expectations of biomonitoring based on eDNA analysis can also be seen in the rapid growth of the number of publications (Rodríguez-Ezpeleta et al., 2021; Tsuji et al., 2019). Moreover, a meta-analysis has shown that 90% of the 63 studies reported the positive relationships between eDNA concentration in a water sample and the abundance and/or biomass of aquatic organisms (Rourke et al., 2022). These results suggest that the eDNA analysis can be further developed and used in the future as a quantitative estimation tool for biodiversity.

Environmental DNA analysis for detecting macroorganisms can be technically categorized into two methods, i.e., species-specific detection and metabarcoding; however, some challenges remain in the quantitative detection of eDNA in both methods (Bylemans et al., 2019). The species-specific detection method using species-specific primers/probe and a real-time PCR system is currently a major eDNA quantitative method. Nevertheless, the development of a species-specific detection system or multiplexing assays of them is time-consuming and costly and requires prior knowledge and assumptions about the species present within a study site (Bylemans et al., 2019; Tsuji et al., 2018; Wozney and Wilson, 2017). Thus, it is unsuitable for the quantitative detection targeting multiple species. Against this background, researchers have recently begun to explore the possibilities of quantitative analysis on eDNA metabarcoding using universal primers targeting taxonomic groups (e.g. Evans et al., 2016; Fraija-Fernández et al., 2020; Kelly et al., 2014; Thomsen et al., 2016). The eDNA metabarcoding enables us to identify community composition including multiple target taxa. However, the number of reads for each taxon output from the high-throughput sequencer is difficult to treat as quantitative information because it can easily vary between samples due to several problems such as PCR inhibitions, primer bias, and library preparation bias (Deiner et al., 2017; Lamb et al., 2019; Lim et al., 2016; Rourke et al., 2022).

Challenge to accurate quantification of eDNA through metabarcoding is an important frontier of eDNA-based biomonitoring, and some approaches have recently been developed (Hoshino et al., 2021; Smets et al., 2016; Ushio et al., 2018). Particularly, the quantitative MiSeq sequencing approach (hereafter, qMiSeq approach) developed by Ushio et al. (2018) allows us to convert the sequence read numbers of detected taxa to DNA copy numbers based on a linear regression between known DNA copy numbers and observed sequence reads of internal standard DNAs. One of the advantages of the qMiSeq approach is that the copy number can be calculated considering the sample-specific effects of PCR inhibition and library preparation bias because a sample-specific standard line is obtained by adding internal standard DNAs to each sample. Accordingly, Ushio et al. (2018) reported the successful quantitative monitoring of eDNA derived from multiple fish species in coastal marine ecosystems by combining the qMiSeq approach and the fish universal primer, MiFish-U (Miya et al., 2015). A further verification experiment also showed a positive correlation between the estimated eDNA concentration and the sound intensity of fish (Sato et al., 2021). These results suggested that the qMiSeq approach is a promising technique for quantitative eDNA-based monitoring of fish community. However, none of the previous studies have been able to compare the observation data with the estimated eDNA concentrations by species and study site. The accumulation of comparative studies on the use of the qMiSeq approach will contribute to the further development and application of quantitative eDNA metabarcoding and maximize its potential as a quantitative monitoring tool for biodiversity (Ushio, 2022).

The objective of this study was to evaluate the performance of the qMiSeq approach as a quantitative monitoring tool for fish community. Here, we compared the qMiSeq approach to the capture-based survey with an electrical shocker at four river systems. Specifically, the followings were tested: 1) for each study site, whether there is a significant positive relationship between the fish eDNA concentrations obtained by the qMiSeq approach and the capture data (fish abundance and biomass), and 2) for each taxon, whether the capture data at each site have significant relationships with on the eDNA concentrations. Based on the results, we discussed the usefulness of the qMiSeq approach for quantitative monitoring of fish community.

## 2 Materials and Methods

### 2.1 Study site and overview of survey design

Field surveys including capture survey and water sampling were conducted in four rivers in western Japan (Fig. 1a, Fig. S1); Yokomichi River (YK; 11th September 2019), Hisakane River (HS; 12th September 2019), Fukuchi River (FK; 8th November 2019) and Ino River (IN; 15th November 2019). The three (HS) or six (YK, FK and IN) study sites were set for each river. Details of each survey site were shown in Table S1. The overview of the survey design is shown in Fig. 1b. Water sampling for eDNA analysis and capture survey with an electrical shocker were performed for each study site.

**Fig. 1.**
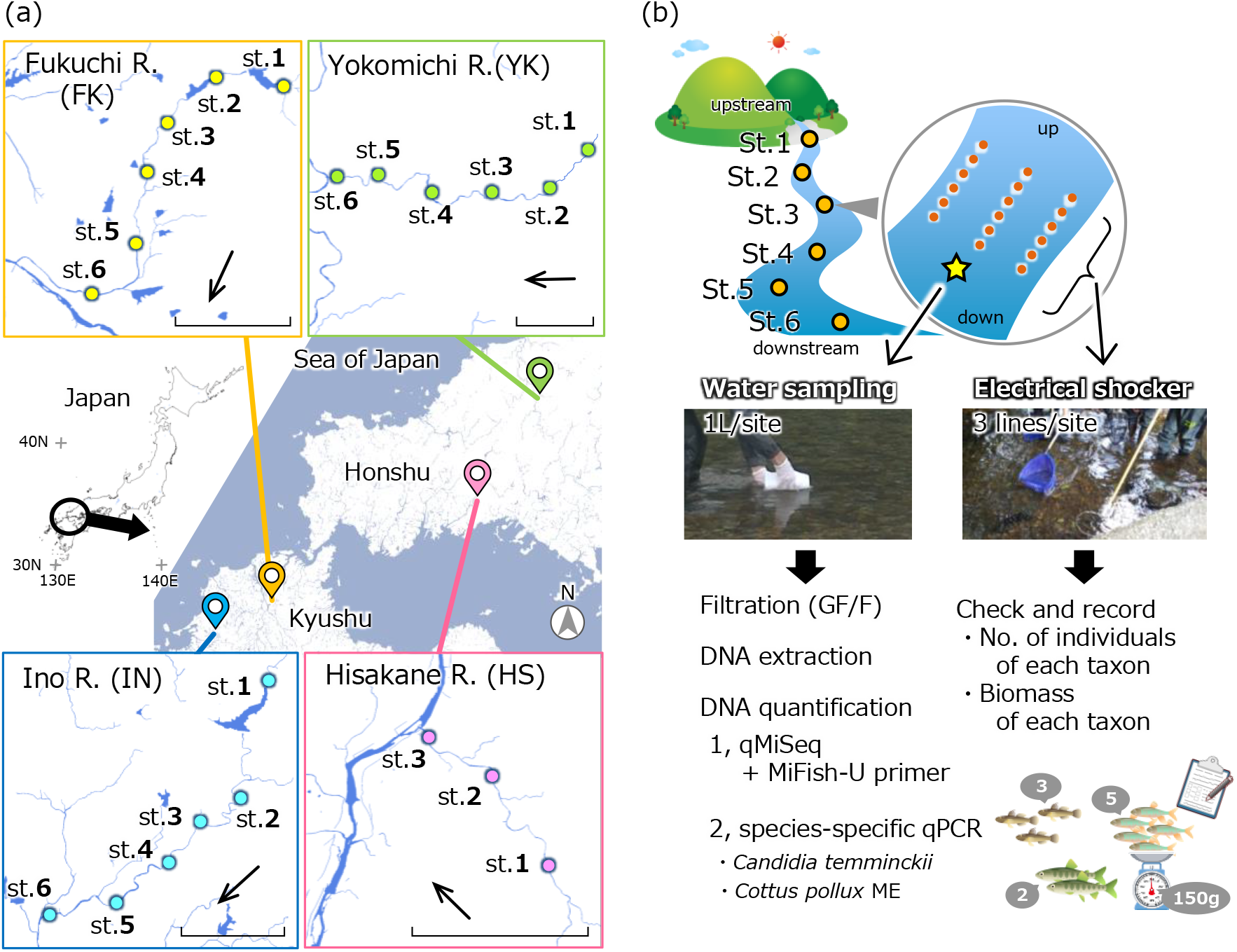
Overview of (a) the study sites and (b) survey design in this study. The bar and arrow in the bottom right-hand corner of the detailed maps in (a) indicate 2 km distance and water flow direction, respectively.

### 2.2 Water sampling and filtration for eDNA analysis and Capture survey by electrical shocker

At each study site, we collected 1 L of surface water using a bleached bottle from the center line of stream at the downstream end of each site (a total of 21 field samples, Fig. 1b). We added benzalkonium chloride (1 mL, 10% w/v; Yamanaka et al., 2017) (Fujifilm Wako Pure Chemical Corporation, Osaka, Japan) to each water sample to preserve eDNA and transported the sample bottles to the laboratory under refrigeration. As a cooler blank for each sampling day, 1 L of deionized water was placed into the same cooler box and thereafter treated in the same way as the collected water samples (a total of four cooler blank samples). The water samples were vacuum filtered using GF/F glass fibre filters (diameter: 47 mm, mesh size: 0.7 μm; GE Healthcare Japan, Tokyo, Japan) within 36 hours after sampling. All filter samples were stored at –20°C until DNA extraction.

After the water sampling, we conducted fish samplings (up to 45 minutes per site) using backpack electroshocker (Smith-Root, LR-20B, USA) and dip nets. Three survey lines were set at each survey site and electric shocks were applied along each line (Fig. 1b). The collected fishes were identified and released within the study reach except for the fishes we could not identify in the field.

### 2.3 DNA extraction from the filter sample

The DNA on each filter sample was extracted following the method described by Environmental DNA Sampling and Experiment Manual ver 2.1 (Minamoto et al., 2021) with minor modifications. First, each filter sample was placed in the upper part of the Salivette tube (SARSTEDT AG & Co. KG, Nümbrecht, Germany). The 220 µL of extraction solution containing 200 μL Buffer AL and 20 µL of proteinase K in DNeasy Blood & Tissue Kit (Qiagen, Hilden, Germany) were added onto each filter and incubated at 56°C for 30 min. After incubation, the Salivette tube was centrifuged at 5,000 × g for 1 min. 220 μL Tris–EDTA (TE) buffer (pH 8.0; Nippon Gene Co., Ltd., Tokyo, Japan) was added onto each filter and re-centrifuged at 5,000 × g for 3 min. After that, 200 μL of ethanol was added to the solution in the bottom part of the Salivette tube and mixed well by gently pipetting. The whole of the mixed solution was transferred to a DNeasy Mini spin column, and the DNA was purified according to the manufacturer’s protocol. The DNA was finally eluted in 100 μL Buffer AE and stored at −20°C.

### 2.4 eDNA quantification: Quantitative metabarcoding with qMiSeq approach

The qMiSeq approach (Ushio et al. 2018) and MiFish-U primers (Miya et al. 2015) was performed to identify fish taxa and quantify their eDNA concentrations simultaneously. The qMiSeq allows us to convert the number of sequence reads into DNA copies without being affected by differences in PCR efficiency by obtaining a sample-specific standard line using internal standard DNAs. The paired-end library preparation with a two-step PCR was performed in 12 µL of reaction mixture according to the described method in Tsuji et al., (2022) (See appendix for details). Briefly, in first-round PCR was performed with MiFish-U-F/R primers (Miya et al. 2015, Table S2), which can amplify fish DNA and three internal standard DNAs (5, 25, 50 copies per reaction, respectively: Table S3). Internal standard DNAs were added to only field samples and cooler blanks. PCR negative control with ultrapure water instead of both eDNA sample and standard DNA mix was added in all first PCR runs. The second-round PCR was performed to add index-sequence and adapter-sequence for the Illumina sequencing platform (Table S2). The indexed products of the second-round PCR were pooled, and the target bands (ca. 370 bp) were excised using 2% E□Gel SizeSelect Agarose Gels (Thermo Fisher Scientific). The prepared DNA libraries were sequenced by 2 × 150 bp paired-end sequencing on the iSeq platform using the iSeq 100 i1 Reagent v1 cartridge (Illumina, CA, USA) with 30% PhiX spike-in.

The bioinformatics analysis was performed using the PMiFish pipeline (Miya et al., 2020; https://github.com/rogotoh/PMiFish). Briefly, first, low-quality tails were trimmed from each sequence read, and the paired-end reads were merged. Then, primer sequences were removed, and identical sequences were merged using UCLUST (USEARCH v10.0.240, Edgar, 2010). The merged sequences with 10 or more reads were assigned to the taxonomy using local BLASTN search with the reference database including all inhabiting freshwater fish taxa around the study sites (MiFish DB ver. 37) and the sequence of used three internal standard DNAs. The top BLAST hits with a sequence identity ≥ 98.5% were applied and used in further analyses. For *Odontobutis obscura* only, sequences detected with sequence identity ≥ 90.0% were also used in the analysis, as they are known to exist in highly genetically differentiated regional populations. As all study sites were in freshwater areas, any saltwater fishes detected were excluded from subsequent analyses. For internal standard DNAs, to obtain a sample-specific standard line, linear regression analysis (lm function in R version 3.6.0 software) was performed using the obtained sequence reads and their known copy numbers (the intercept was set as zero). For each sample, the number of eDNA copies per litre for each taxon was calculated as qMiSeq eDNA concentration using each sample-specific standard line: qMiSeq eDNA concentration = the number of iSeq sequence reads/regression slope of the sample-specific standard line (Table S4).

### 2.5 eDNA quantification: real-time quantitative PCR (qPCR) for two taxa

The real-time qPCR for *Candidia temminckii* and *Cottus pollux* ME were performed using a StepOne-Plus Real-Time PCR system (Applied Biosystems, FosterCity, CA, USA). The eDNA samples from the Yokomichi River were omitted from the comparison examination because they were used in another study and there was no remaining volume for analysis. Species-specific primers for each target species were developed in this study (Table S2), and their specificity was tested by in silico and in vitro tests. All qPCR reactions were performed in a total 15 µL volume and triplicated (See appendix for details of real-time qPCR conditions and species-specific primer development).

### 2.6 Statistical analyses

For the following analyses, the R ver. 3.6.0 software was used (R Core Team. R, 2019). The significance level was set at 0.05 in all analyses. To examine whether the DNA concentrations quantified by the qMiSeq approach reflect those quantified by species-specific qPCR, the eDNA concentrations of *C. temminckii* and *C. pollux* ME quantified by the two methods were compared using the linear regression analysis (lm function in R). The differences in fish community compositions among sites were visualised using nonmetric multidimensional scaling (NMDS) with 500 permutations and cluster analysis with group average method. The Bray-Curtis dissimilarity index was used as the fish community dissimilarity in both NMDS and k-means clustering to use the DNA copy number as quantitative information. Based on k-means clustering, a total of 21 study sites were divided into five clusters (see Results). The number of cluster, five, was determined on the basis of Calinski criterion values (Fig. S2). The differences in eDNA release rate and capture efficiency are expected between study sites with largely different fish community compositions due to differences in geographical distribution, ecology, and behavior of each species, which could distort the relationship between observed DNA concentrations and capture data. Thus, qMiSeq eDNA concentrations and capture data (abundance and biomass of each taxon) were compared using the Kendall-rank correlation test by classifying the data in two patterns as follows: (1) using data from all survey sites and (2) using each cluster site data classified by similarity of fish community. Moreover, for eleven taxa which were detected from three and over sites by each of qMiSeq approach and electrical shocker among all study sites, the pooled dataset of qMiSeq eDNA concentrations and capture data (abundance or biomass at each survey site) was created for each taxa. For each taxa, qMiSeq eDNA concentrations and capture data were compared using generalized linear models (GLM; the glm.nb function in the MASS package in R, Venables and Ripley, 2002) assuming that eDNA copy numbers follow a negative binomial distribution with log-link function (Table 1). Here we have chosen GLMs as a number of previous studies have suggested that the relationship between abundance or biomass of each fish species and eDNA concentration is non-linear (e.g. Coulter et al., 2019; Doi et al., 2017). Additionally, the eleven taxa were divided into four benthic fish taxa group and seven pelagic fish taxa group, and the same analysis was carried out for each group as described above. Note, however, that data on *O. obscura* and *O. hikimius*, which are closely related and have almost identical ecology, were merged and treated as a single taxon.

**Table 1.** Results of GLM for relationships between qMiSeq eDNA concentration (copies/L) and abundance or biomass for each taxon. The scientific name basically followed Nakabo (2013), but *Cottus* sp. were denoted as *Cottus pollux* ME because it was known that *Cottus* sp. in the study sites have mitochondria sequences consistent with *Cottus pollux* ME regardless of morphology (Kanno et al., 2018).

| Taxon | No. of individuals |  |  |  | Biomass (g) |  |  |  |
| --- | --- | --- | --- | --- | --- | --- | --- | --- |
|  | Estimate | SE | z | P | Estimate | SE | z | P |
| Four benthic fish taxa | 0.085 | 0.018 | 4.86 | < **0.0001** | 0.015 | 0.003 | 5.39 | < **0.0001** |
| *Odontobutis obscura* and *O. hikimius* | 0.082 | 0.022 | 3.67 | < **0.001** | 0.012 | 0.004 | 3.13 | < **0.01** |
| *Tridentiger* spp. | 0.034 | 0.023 | 1.51 | 0.13 | 0.012 | 0.008 | 1.46 | 0.14 |
| *Cottus pollux* ME | 0.073 | 0.018 | 4.04 | < **0.0001** | 0.016 | 0.008 | 1.92 | 0.06 |
| *Pseudogobio esocinus esocinus* | -0.117 | 0.097 | -1.21 | 0.23 | -0.005 | 0.004 | -1.29 | 0.20 |
| Seven pelagic fish taxa | 0.030 | 0.005 | 5.74 | < **0.0001** | 0.005 | 0.001 | 4.50 | < **0.0001** |
| *Carrassius* spp. | 0.039 | 0.028 | 1.42 | 0.16 | 0.002 | 0.001 | 1.57 | 0.12 |
| *Candidia temminckii* | 0.011 | 0.006 | 2.02 | < **0.05** | 0.004 | 0.002 | 2.61 | < **0.01** |
| *Pungtungia herzi* | 0.033 | 0.019 | 1.71 | 0.09 | 0.008 | 0.004 | 2.21 | < **0.05** |
| *Opsariichthys platypus* | 0.040 | 0.000 | 88.77 | < **0.0001** | 0.010 | 0.008 | 1.29 | 0.20 |
| *Salvelinus* sp. | 0.474 | 0.099 | 4.79 | < **0.0001** | 0.018 | 0.005 | 3.61 | < **0.0001** |
| *Oryzias latipes* | 0.209 | 0.156 | 1.34 | 0.18 | 1.458 | 0.952 | 1.53 | 0.13 |
| *Rhynchocypris oxycephalus jouyi* | 0.099 | 0.033 | 3.01 | < **0.01** | 0.011 | 0.004 | 2.62 | < **0.01** |

## 3. Results

### 3.1 Overview of qMiSeq approach results and the comparison of eDNA concentrations quantified by the qMiSeq approach with those estimated by qPCR

The iSeq paired-end sequencing (2□×□150 bp) for the 27 libraries (21 field samples, four cooler blanks, two PCR negative controls) yielded a total of 4.71 million reads (Q30 = 95.2; PF = 67.3 %). Negligible sequence reads were detected from PCR negative controls (maximum 75 reads; Table S5), which were ignored in the subsequence analyses. The sequence reads of internal standard DNAs in field samples and cooler blanks had a significant positive relationship with the copy numbers (lm, P < 0.01, R^2^ > 0.93; Table S4). The coefficients of the linear regressions varied from 174.1 (IN st. 4) to 900.8 (IN cooler blank) (Table S4), suggesting that sequence reads are proportional to the DNA copy numbers in a single sample. Thus, we converted sequence reads of detected taxa in each sample to the DNA copies using the coefficient of the `sample-specific regression line. In the comparison of DNA concentrations of the three taxa calculated by qMiSeq approach and species-specific qPCR, a significant positive relationship was found for each taxa (linear regression analysis; P < 0.001, R^2^ = 0.81, estimate 0.32 for *C. temminckii*; P < 0.001, R^2^ = 0.99, estimate 0.58 for *C. pollux* ME; Fig. 2).

**Fig. 2.**
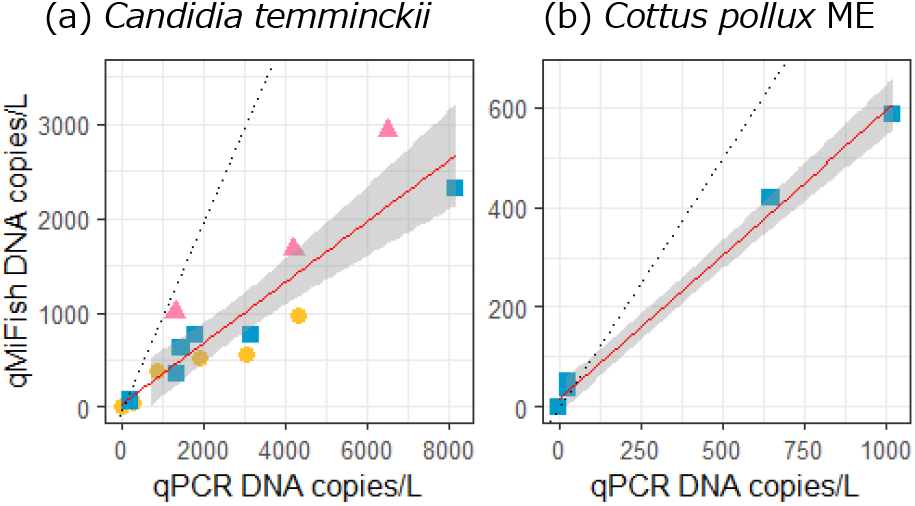
Relationship between the eDNA concentration quantified by qMiSeq approach and that by species-specific qPCR for (a) *C. temminckii* and (b) *C. pollux* ME. Yellow circles (FK), pink triangles (HS) and blue squares (IN) indicate each study river, respectively. The eDNA samples from the Yokomichi River were omitted from the comparison examination because they were used in another study and there was no remaining volume for analysis. Solid and dashed lines indicate linear regression lines (all regression lines are significant, P < 0.001) and 1:1 line, respectively. The shaded area around each linear regression lines corresponds to the 95% confidence intervals.

### 3.2 Comparison of species richness observed by qMiSeq and electrical shocker

The qMiSeq approach consistently detected more species than the capture-based survey using an electrical shocker (Fig. 3). On the other hand, one or two taxa that were detected by the electrical shocker were not detected by the qMiSeq approach (i.e., false negative) in several study sites: HS st.3, *Liobagrus reini*; FK st. 4, *Opsariichthys platypus* and *Oryzias latipes*; FK st. 5, *Cobitis matsubarae*; FK st.6, *Squalidus gracilis gracilis* and *Tachysurus nudiceps*; IN st. 5; *S. gracilis gracilis*. The false negative result in qMiSeq results for *Cobitis matsubarae* at FK st. 5 was excluded from all subsequent analyses as it was likely due to a lack of reference DNA database (see Discussion for detail). On the other hand, no false negatives were observed at 16 out of a total of 21 sites. At almost all sites, *Rhinogobius* spp. was detected by both or either of methods, but it was excluded from subsequent analyses because they contain many unknown haplotypes and many closely related species that cannot be discriminated and/or detected based on the target region of MiFish primer. Also, at IN st. 5, *Cyprinus carpio* were detected by both methods, but were excluded from subsequent analyses because five individuals (approximately 50 cm, total length) were also visually identified immediately upstream of the study section. Their eDNA almost certainly flowed into the study section and was considered to be noise in this study. The fish taxa detected only by qMiSeq approach were the taxon collected from other sites in the same river, a rare native species likely to be inhabited or an invasive species in the early stages of invasion (Table S5).

**Fig. 3.**
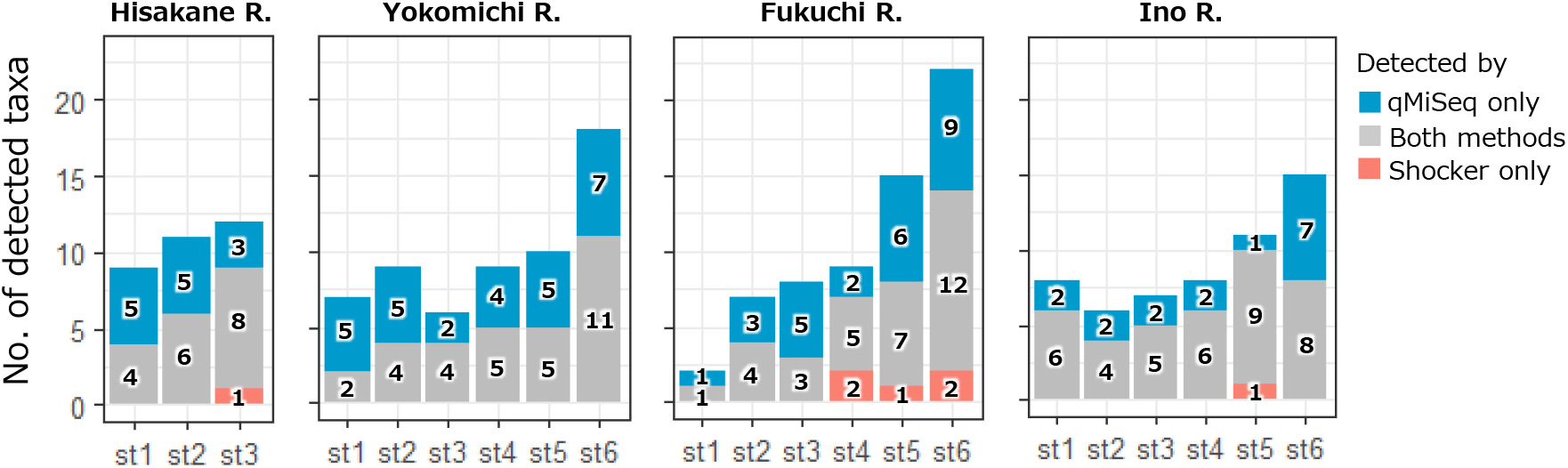
The number of taxa detected by eDNA metabarcoding with qMiSeq (blue), a capture-based survey using electrical shocker (pink) or both methods (grey) at each study site of each river.

### 3.3 Fish community structure

The dissimilarity of the fish community structure among study sites was shown in the nonmetric multidimensional scaling (NMDS) biplot (Fig. 4a). A total of 20 research sites (FK st. 1 was excluded due to absebce of detected taxa) were categorised into five clusters by k-means clustering. The clusters tended to be divided by the down and up stream. Cluster 3 and 4 consisted only of the study sites at Yokomichi River.

**Fig. 4.**
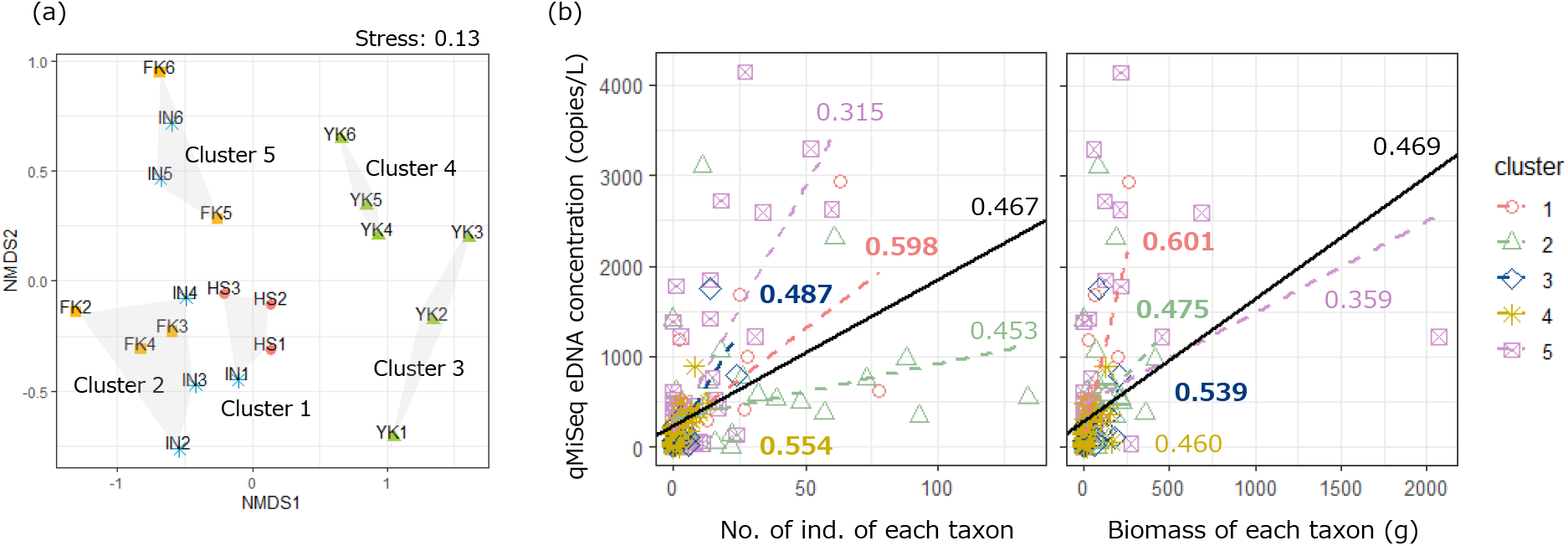
Visualisation of fish communities detected by qMiSeq approach at each study site: (a) Nonmetric multidimensional scaling (NMDS) ordination plot of fish communities and (b) relationship between the qMiSeq eDNA concentration and the abundance or biomass of each taxon for all study sites and study sites included in each site cluster. NMDS ordination plot and site clustering in (a) panel were based on Bray-Curtis dissimilarity index and k-means clustering. Solid black line in (b) panels indicates a linear regression line obtained using all study sites data. Coloured dashed lines indicate linear regression lines obtained using each cluster site data. Coloured letters next to regression lines indicate the rank correlation coefficients (Kendall’s τ). If the Kendall’s τ value obtained using data from each cluster site was greater than that obtained using data from all study sites, the rank correlation coefficients were expressed in bold.

### 3.4 Comparison of the qMiSeq eDNA concentrations with the capture data by electrical shocker

When all sites were used for analysis, both the number and biomass of captured fishes had significant positive correlations with the qMiSeq eDNA concentrations (Kendall rank correlation test, Ps < 0.0001, τ = 0.467 for abundance and τ = 0.469 for biomass; Fig. 4b, Table S6). Additionally, the significant positive correlations were also found for all site clusters (Kendall rank correlation test, both the number and biomass, P < 0.01 for all site clusters; Fig. 4b, Table S6). The Kendall’s τ value varied widely among clusters (0.315 to 0.598 for abundance; 0.359 to 0.601 for biomass). Moreover, for seven out of eleven taxa which were detected from three and more sites by each of qMiSeq approach and electrical shocker, we found significant positive relationships between the qMiSeq eDNA copy numbers at each site and the number of captured individuals and/or the total biomass (GLM; Fig. 5, Table 1). When eleven taxa were grouped into four benthic fish taxa and seven pelagic fish taxa, there were significant positive relationships between the qMiSeq eDNA concentration and both the number of captured individuals and the total biomass (GLM; Table 1).

**Fig. 5.**
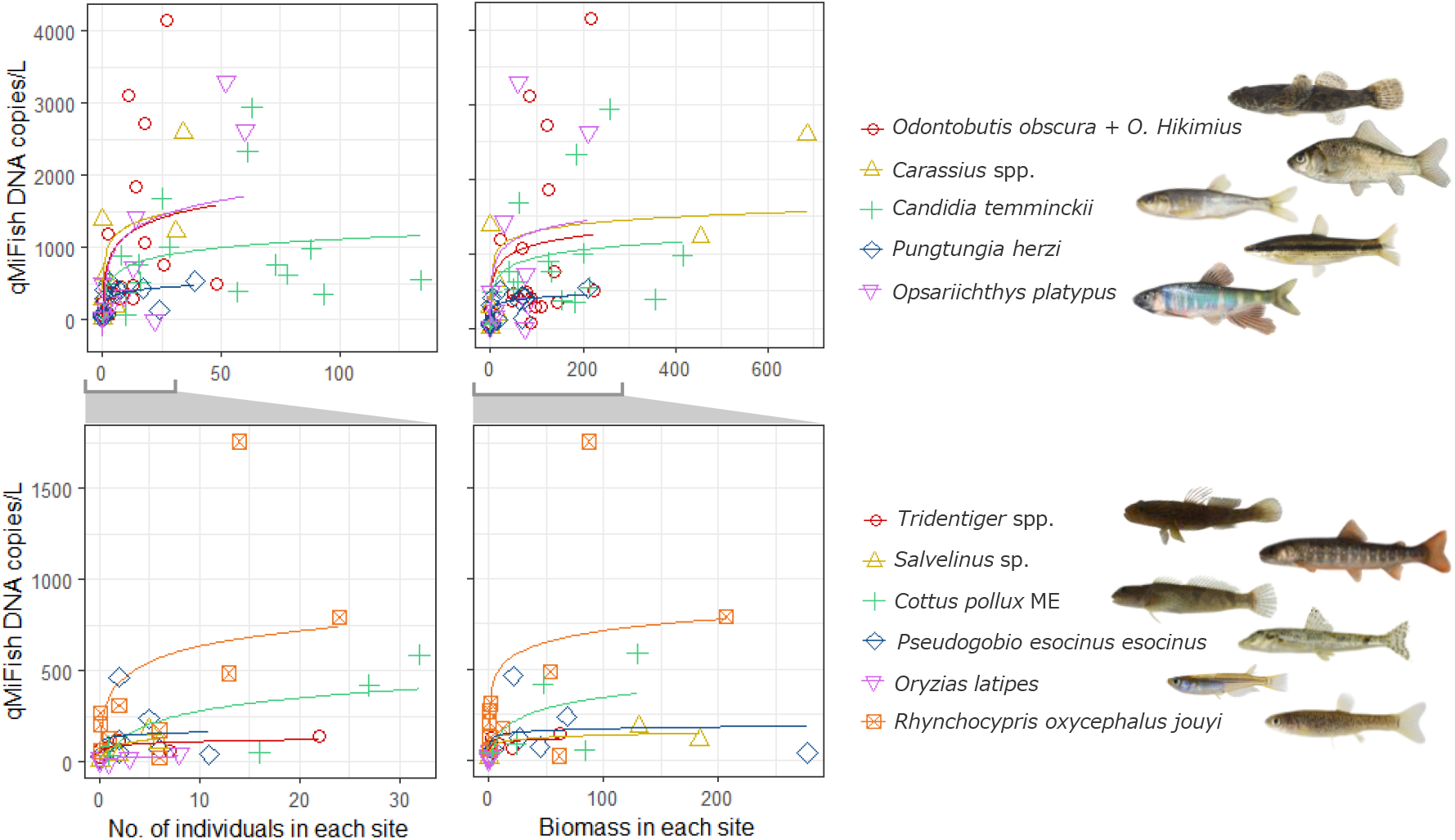
Relationship between the qMiSeq eDNA concentrations quantified and capture data (abundance or biomass) for each taxon at each study site. Each taxa was allocated to either upper or lower panels according to the abundance/biomass level. Each coloured solid line indicates nonlinear regressions of a generalized linear model (GLM) based on negative binomial distribution. Photo copyright: *O. obscura, C. temminckii, P herzi, C. pollux* ME, *P. esocinus esocinus* and *R. oxycephalus jouyi* for Mr. S. Kunumatsu; *Carassius* spp. and *O. latipes* for Mr. Y. Fuke; *O. platypus* and *Salvenius* sp. for ffish.asia (https://ffish.asia/, 2022.04.18 downloaded); *Tridentiger* spp. for S.T.

## 4. Discussion

### 4.1 eDNA metabarcoding detected higher species richness than electrofishing survey

We found that eDNA metabarcoding with the qMiSeq approach could detect not only almost all the taxa captured using the electric shocker but also other taxa that could be false negatives in the capture surveys. This result is consistent with several previous studies that have reported advantages of eDNA metabarcoding compared with traditional capture-based methods in terms of detection sensitivity on biodiversity assessment (Andruszkiewicz et al., 2017; Bylemans et al., 2018; Civade et al., 2016; Deiner et al., 2016; Hänfling et al., 2016; Miya et al., 2015; Nakagawa et al., 2018; Sato et al., 2017; Shaw et al., 2016; Thomsen et al., 2012; Valentini et al., 2016; Yamamoto et al., 2017). On the other hand, we should always be aware the possibility of false negative. In this study, although almost all taxa were detected, a total of six taxa were missed by eDNA metabarcoding at the sampling site level. Three main causes of false negative results have been suggested in eDNA metabarcoding: (1) low concentration of target DNA, (2) PCR drop-out because of primer mismatch and (3) incompleteness and inaccuracy of the reference sequence database (Evans et al., 2017; Jane et al., 2015; Miya et al., 2020).

### 4.2 The significant positive correlation between eDNA concentrations quantified by qMiSeq and qPCR

The significant positive relationship between eDNA concentrations quantified by the qMiSeq approach and species-specific qPCR indicated that the qMiSeq approach successfully estimated the number of sequence reads of each taxon to the eDNA copy number. For both taxa considered, the eDNA copy numbers calculated by the qMiSeq approach tended to be smaller than those estimated by qPCR. This observation was consistent with those reported in the previous study (Ushio et al., 2018). Additionally, eDNA metabarcoding has been recognised to be less sensitive than species-specific detection due to the effects of technical issues such as primer bias, PCR bias and the potential reduced amplification efficiency in tailed-PCR (Bylemans et al., 2019; Harper et al., 2018; Nichols et al., 2018). However, this characteristic would not be a serious problem when we compare the eDNA concentrations quantified by the qMiSeq approach on a species-by-species basis.

### 4.3 Calculated eDNA concentration by qMiSeq reflected both fish abundance and biomass in each study site

It was worthy of note that both abundance and biomass of inhabiting fishes had positive correlations with the qMiSeq eDNA concentrations. This study is the first to demonstrate the relationships between the quantitative capture data of fishes and their eDNA concentrations quantified using eDNA metabarcoding. Meanwhile, Kendall’s rank correlation coefficients were low in both abundance and biomass (τ < 0.5), possibly because differences in eDNA release and/or degradation rates among taxa inhabiting the study sites, capture efficiency with electric shockers, and environmental conditions could affect the results.

The rank correlation coefficients (Kendall’s τ) tended to be improved by dividing the samples into site clusters based on the dissimilarity of fish community compositions, compared to when all study sites were considered. The results suggested that the study site clustering based on fish community compositions has mitigated the effects of the differences of the eDNA ecology and/or dynamics such as release rate and/or degradation rates for each taxon and dispersion processes on the relationship between qMiSeq eDNA concentration and fish capture data. In contrast, at some clusters, Kendall’s τ values were decreased. Particularly, in cluster 5 sites, Kendall’s τ values decreased for both abundance and biomass and were smallest in all site clusters. The cluster 5 consisted only of downstream sites (IN st. 5, 6 and FK st. 5, 6), and vegetation covered large areas of the channel, mainly along the river bank (Figure S1). Relatedly, the taxa with highly biased relationships between qMiSeq eDNA concentrations and capture data had ecological characteristics such as a preference for vegetation (*Pungtungia herzi*) and rock crevices (*O. obscura*), hiding in the sandy bottom (*Pseudogobio esocinus esocinus*) and high swimming ability (*C. temminckii*) (Hosoya, 2019; Nakabo, 2013). The environmental condition and these ecological characteristics would have strongly influenced capture efficiency in the electrical shocker survey. Taxa hiding in vegetation, rock crevices and sandy bottom were difficult to capture because electric shocks cannot reach them, or they faint in unseen crevices. Fish taxa with high swimming ability were able to quickly escape from the investigator’s position or hide under vegetation. In this study, it was difficult to conduct an exhaustive fish capture by closing the study site section due to the limitation of labour, manpower and time for the survey. These facts may indicate that differences in the capture efficiency of the electrical shocker survey have influenced the results of this study, although the magnitude of the impact could depend on the environmental conditions and the ecological characteristics of taxa that inhabit at each study site.

### 4.4 qMiSeq eDNA concentration reflected both fish abundance and biomass in each detected taxon

For seven taxa out of eleven taxa with higher detection frequency in both methods, we demonstrated that the qMiSeq eDNA concentration was significantly related to the abundance and/or biomass. Additionally, qMiSeq eDNA concentration and both the number of captured individuals and total biomass were significantly related when the eleven taxa were grouped into benthic or pelagic fish taxa. It was consistent with the results of many previous studies showing positive relationships between eDNA concentration and the abundance and/or biomass of target species (90% of relevant 63 papers published by 2020, Rourke et al., 2022). This observation supports the potential and usefulness of the qMiSeq approach as a quantitative monitoring tool of biodiversity beyond simply identifying species presence. Our result in this study was significant for two reasons.

First, to the best of our knowledge, there have been no studies comparing eDNA concentrations of various macroorganism taxa obtained by quantitative metabarcoding methods, such as the qMiSeq approach, with quantitative capture data for each taxon. To the future implementation of quantitative eDNA metabarcoding methods for quantitative biodiversity monitoring, we should continue the detailed and repeated research efforts to determine whether the results obtained from conventional survey methods and eDNA-based methods yield similar measurements for the taxa of interest (Deiner et al., 2017). We believe that this study was the first step in this process. Second, it is worth mentioning that the comparisons of eDNA concentrations and capture data in this study have been performed using pooled eDNA concentration data for each species derived from multiple different samples. In general, the direct comparison of sequence reads count among different samples to quantitatively estimate biodiversity lead to erroneous conclusions because the number of sequence reads will vary among samples depending on the effects of PCR inhibition, primer bias, library preparation bias and sequencing depth, etc (Hoshino et al., 2021; Ushio et al., 2018). In most cases, researchers used the proportional abundance of sequence read counts in a sample for each detected taxon instead of a direct comparison of their sequence read counts (e.g. Fraija-Fernández et al., 2020; Goutte et al., 2020; Thomsen et al., 2016). However, as differences in the number of taxa detected per sample inherently bias the proportion of sequence read counts (i.e. an increase in the total number of species would reduce the proportional abundance), it is difficult to apply to quantitative comparisons of target taxa among sites, seasons and years with different community compositions (Bylemans et al., 2019). In contrast, although the qMiSeq approach also cannot mitigate the effects of primer bias (see below), it allows avoiding the effects of PCR inhibition, library preparation bias and sequencing depth (Ushio et al., 2018). We reasonably compared the eDNA concentrations of each taxon from multiple samples and demonstrated the usefulness of the analytical advantages of the qMiSeq method for quantitative eDNA metabarcoding for macroorganisms.

Using the qMiSeq approach, although a significant relationship was observed between qMiSeq eDNA concentration and capture data for 7 taxa and benthic or pelagic fish taxa group, the obtained values varied widely between and within the taxon. It is obvious that the relationship between qMiSeq eDNA concentration and abundance and/or biomass of target taxon observed by field experiments can be obscured by biological and non-biological factors, which is the limitation of analytical method and survey design (Deiner et al., 2017; Rourke et al., 2022). In this study, the environmental conditions such as water temperatures and flow velocity at the time of water sampling varied widely among study sites (Table S1), and there may have been associated differences in the characteristics of the ‘ecology of eDNA’ within each taxon (e.g., its origin, state, fate and transport) (Barnes and Turner, 2016; Collins et al., 2019; Jane et al., 2015; Tsuji et al., 2017). Additionally, as mentioned above, the limitations of capture survey using electrical shocker was likely to have caused significant fluctuations in the observed values.

Furthermore, we also need to mention the effect of primer bias, which is an analytical limitation of eDNA metabarcoding. Primer bias is recognised as one of the main causes of the distortion of the relative abundance of amplified DNA in a sample in DNA metabarcoding data (Elbrecht and Leese, 2015; Lamb et al., 2019; Nester et al., 2020; Piñol et al., 2015). The MiFish-U primers used in this study has been proven to have high primer universality by many previous studies (e.g. Collins et al., 2019; Miya et al., 2020; Zhang et al., 2020). Thus, we believe that the distortion of results due to primer bias was minimised in this study.

## Conclusions

We here compared the eDNA concentrations quantified by the qMiSeq approach and the results of the capture-based survey using an electrical shocker to evaluate the performance of the qMiSeq approach as a quantitative monitoring tool for fish community. We found positive relationships between the qMiSeq eDNA concentrations and both the abundance and biomass of each captured taxon. Together, our results suggested that eDNA metabarcoding with the qMiSeq approach is a suitable and useful tool for quantitative monitoring of fish community. The simplicity of eDNA analysis will reduce barriers forassessing changes in species abundance, evenness and richness and will facilitate the collection of valuable information for better understanding biodiversity changes.

## Authors’ contributions

S.T., R.I. and Y.A. conceived and designed research. R.I, S.M. M.S. and T.K. performed a field survey. S.T. and R.N. performed molecular experiments and data analysis. S.T. wrote the early draft and completed it with significant inputs from all authors.

## Data availability

Full details of the eDNA metabarcoding with qMiSeq approach and species-specific qPCR results are available in the supporting information (Table S5 and S7). All raw sequences were deposited in the DDBJ Sequence Read Archive (accession number: DRA013858).

## Acknowledgements

We thank laboratory members of Akamatsu laboratory, Yamaguchi University, and Inui Laboratory, Fukuoka Institute of Technology University, for help with the field sampling. We thank Mr. S. Kunimatsu and Mr. Y. Fuke for providing fish pictures. We thank Dr. M. Ushio for his valuable comments and advices on the draft. This is supported by the River Fund of The River Foundation, Japan (2019-5211-030). This work was supported by YU Project for Formation of the Core Research Center.

---

**Fig. S1.**
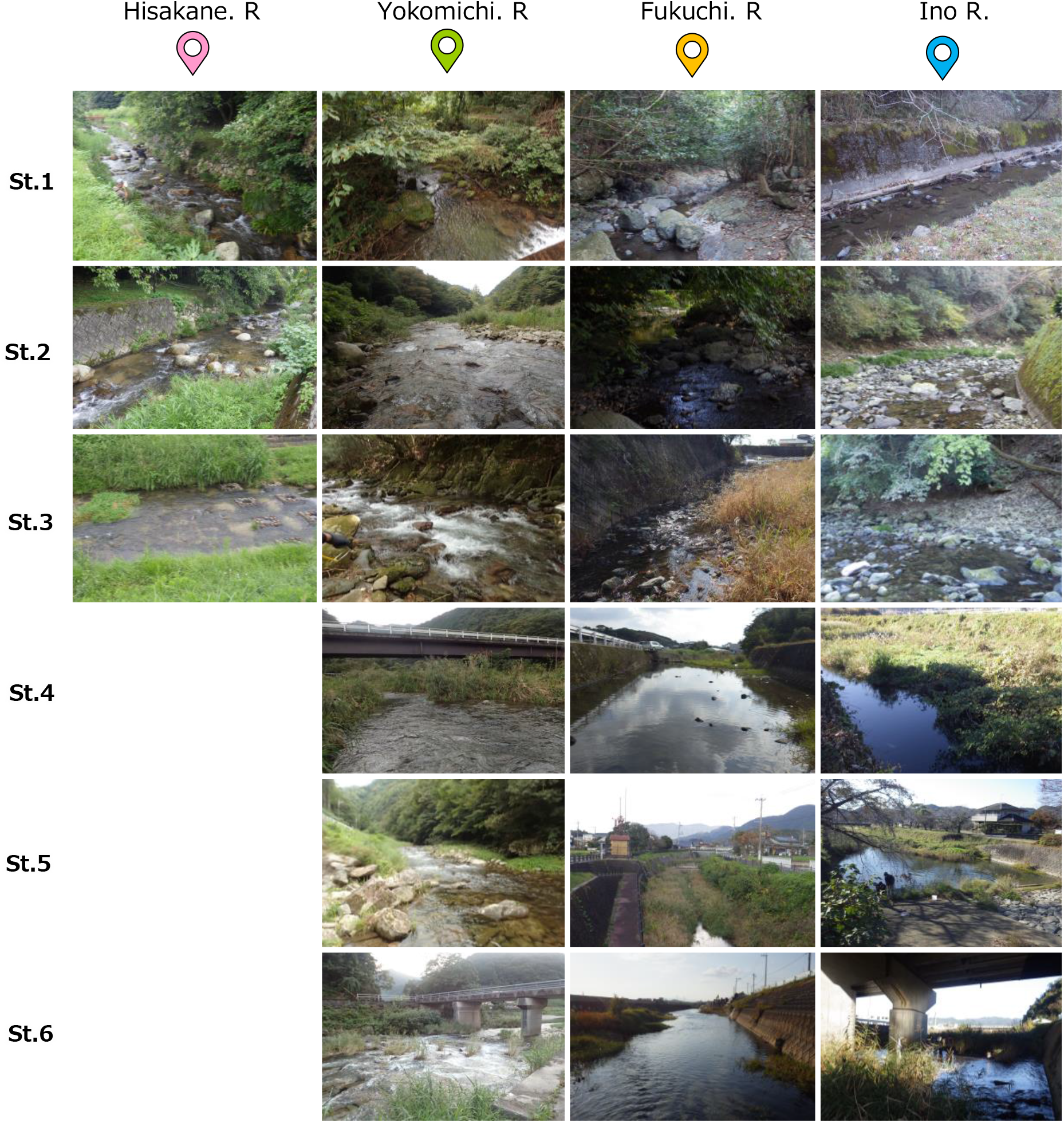
Picture of each survey site.

**Fig. S2.**
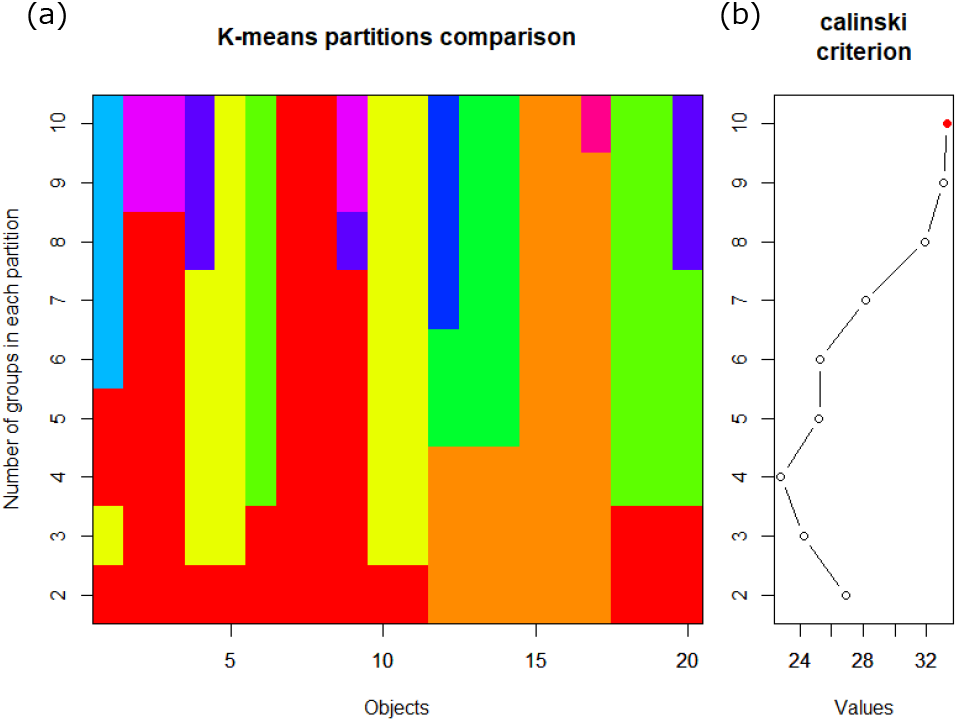
(a) K-means partitions comparison and (b) Calinski values.

