## Appendix for "Quantitative environmental DNA metabarcoding shows high potential as a novel approach to quantitatively assess fish community"

**Appendix: Further Methodological Details**

***Paired-end library preparation in qMiSeq approach***

Quantitative eDNA metabarcoding (qMiSeq approach; Ushio et al. 2018) was performed using a fish universal primer set, MiFish-U primers (Miya et al. 2015), and three standard DNAs. The sequences of MiFish primers (MiFish-U-F/R) were shown in Table S2. The first‐round PCR (first PCR) was performed in a 12-µL total volume containing 6.0 µL of 2× KAPA HiFi HotStart ReadyMix (KAPA Biosystems, MA, USA), 0.72 µL each MiFish-U primer (5 µM), 2.56 µL sterilised distilled H_2_O, 1.0 µL standard DNA mix (number of copies of each standard DNA in 1 µL; 5 copies for std. 1, 25 copies for std. 2, 50 copies for std.3: Table S3) and 1.0 µL eDNA sample. The thermal conditions for the first PCR are as follows: 3 min at 95°C, 40 cycles of 20 s at 98°C, 15 s at 65°C, 15 s at 72°C and 5 min at 72°C. PCR negative control with ultrapure water instead of both eDNA sample and standard DNA mix was added in all first PCR runs. First PCR was performed in eight replicates for each eDNA sample and the negative control. To remove primer dimer and dNTPs, the pooled eight replicates of PCR products were purified using GeneRead Size Selection Kit (Qiagen, Hilden, Germany) accordingly to manufacturer instructions. The purified first PCR products were diluted to 0.1 ng/µL and used as the second-round PCR template (second PCR).

The second PCR was performed in a 12-µL total volume containing 6.0 µL of 2× KAPA HiFi HotStart ReadyMix, 2.0 µL of each index primer (1.8 µM), 1.0 µL sterilised distilled H_2_O, 1.0 µL of the purified first PCR product (0.1 ng/µL). The sequences of the second PCR primers with indexes (*X*) were shown in Table S2. Different combinations of indexing primers were used to distinguish output data in parallel sequencing. The thermal conditions for the second PCR are as follows: 3 min at 95°C, 12 cycles of 20 s at 98°C, 15 s at 72°C and 5 min at 72°C. The indexed products of the second PCR were pooled, and the target bands (ca. 370 bp) excised using 2% E‐Gel SizeSelect Agarose Gels (Thermo Fisher Scientific). DNA concentrations were adjusted to 30 pM (assuming 1 bp DNA has a molecular weight of 660 g/mol) and sequenced on the iSeq platform using an iSeq 100 i1 Reagent v1 cartridge (Illumina, CA, USA) with 30% PhiX spike‐in. All raw sequences obtained in the present study were deposited in the DDBJ Sequence Read Archive (accession number: DRA013858).

***Species-specific primer development for three species***

We developed species-specific primers for *Candidia temminckii*, and *Cottus pollux* ME. For each target species, mitochondrial DNA cytochrome *b* (cyt *b*) sequence data was downloaded from the National Center for Biotechnology Information (NCBI) database (https://www.ncbi.nlm.nih.gov/). Also, for the closely related species for each target species, cyt *b* sequences were downloaded with same manner. The species considered as closely related species for each target species were listed in Table S8. Species-specific primers for each target species were manually designed to meet either or both of the following criteria: (1) mismatches with closely related species within two base pairs from the 3′ ends of the primer and (2) there are four or more mismatches with closely related species throughout the design site of the primer sequence. The specificities of the designed primer for each target species were checked by in silico test using Primer-BLAST (http://www.ncbi.nlm.nih.gov/ tools/primer-blast/) with default settings. We confirmed that each primer sets designed for *C. temminckii*, and *C. pollux* ME would not amplify the DNA of non-target species inhabiting the western Japan. Additionally, for *C. temminckii* and *C. pollux* ME, the TaqMan probe was designed to contain at least one or more base mismatch with a non-target species within the amplified region of the developed species-specific primers.

Moreover, the in vitro test was performed using extracted genomic DNA from each target species (for three individuals each) and non-target species (for one or three individuals each) (Table S8). In all PCR and real-time PCR plates, three replicates were used for PCR negative controls to assess the occurrence of cross-contamination.

First, DNA amplification tests by PCR and electrophoresis were performed. Genomic DNA was amplified with three PCR replicates for each sample using a Mastercycler nexus System (Eppendorf, Hamburg, Germany). Each PCR reaction mixture (15 μL total volume) contained 900 nM of each primer (forward and reverse), 0.075 μL AmpErase Uracil N-Glycosylase (Thermo Scientific), and 2 μL of the DNA sample (20 pg) in 1 × PCR master mix (TaqMan Environmental Master Mix 2.0; Thermo Scientific). The PCR thermal conditions were set as same manner with real-time qPCR (2 min at 50°C and 10 min at 95˚C, 55 cycles of 15s at 95°C and 1 min at 60°C). The species-specific amplification for each primer set was confirmed by electrophoresis of PCR products. Next, DNA amplification tests by real-time PCR with primer probe set was performed. Genomic DNA was amplified with three PCR replicates for each sample using a StepOnePlus Real-Time PCR System (Thermo Fisher Scientific, Waltham, MA). The composition of PCR reaction mixture and PCR thermal conditions were all the same as in ‘*Environmental DNA detection: real-time qPCR with TaqMan probe method*’(described below). The species-specific amplification for each primer probe set was confirmed.

***Environmental DNA detection: real-time qPCR with TaqMan probe method***

The reaction mixture consisting of 900 nM of forward and reverse primer, 125 nM of TaqMan probe, 0.075 μL AmpErase Uracil N-Glycosylase (Thermo Fisher Scientific, MA, USA) and 2 µL DNA template in 1 ×  TaqMan Environmental Master Mix (Thermo Fisher Scientific). For each qPCR plate, the 2 μL of ultrapure water and dilution series of commercially synthesized artificial DNA fragments including each target species (3.0 × 10^1^–3.0 × 10^4^ copies per reaction) were simultaneously analysed as a PCR negative control or standard sample. All real samples, negative controls and standards were analysed in triplicate. The thermal conditions were 2 min at 50°C and 10 min at 95˚C, 55 cycles of 15s at 95°C and 1 min at 60°C.
